# *In-silico* analysis of SARS-CoV-2 genomes: Insights from SARS encoded non-coding RNAs

**DOI:** 10.1101/2020.03.31.018499

**Authors:** Neha Periwal, Sankritya Sarma, Pooja Arora, Vikas Sood

**Affiliations:** Department of Biochemistry, Jamia Hamdard, New Delhi- 110062, India; Department of Zoology, Hansraj College, University of Delhi, North Campus, Delhi-110007, India

**Keywords:** Coronavirus, SARS-CoV-2, miRNAs, Non-coding RNAs

## Abstract

Recently a novel coronavirus (SARS-CoV-2) emerged from Wuhan, China and has infected more than 571000 people leading to more than 26000 deaths. Since SARS-CoV-2 genome sequences show similarity with those of SARS, we sought to analyze all the available SARS-CoV-2 genomes based on the insights obtained from SARS genome specifically focusing on non-coding RNAs. Here, results are presented from the dual approach i.e identifying host encoded miRNAs that might regulate viral pathogenesis as well as identifying viral encoded miRNAs that might regulate host cell signaling pathways and aid in viral pathogenesis. Analysis utilizing first approach resulted in the identification of 10 host encoded miRNAs that could target the genome of both the viruses (SARS-CoV-2 and SARS reference genome). Interestingly our analysis revealed that there is significantly higher number of host miRNAs that could target SARS-CoV-2 genome as compared to the SARS reference genome. Results from second approach involving SARS-CoV-2 and SARS reference genome identified a set of virus encoded miRNAs which might regulate host signaling pathways. Our analysis further identified a similar “GA” rich motif in SARS-CoV-2 genome that was shown to play a vital role in lung pathogenesis during severe SARS infections. Hence, we successfully identified human and virus encoded miRNAs that might regulate pathogenesis of both these coronaviruses and the fact that more number of host miRNAs could target SARS-CoV-2 genomes possibly reveal as to why this virus follows mild pathogenesis in healthy individuals. We identified non-coding sequences in SARS-CoV-2 genomes that were earlier reported to contribute towards SARS pathology. The study provides insights into the overlapping sequences among these viruses for their effective inhibition as well as identifying new drug targets that could be used for development of new antivirals.

## Introduction

Coronaviruses are the group of enveloped viruses with positive-sense single stranded RNA as the genetic material. These viruses cause mild to severe respiratory and intestinal infections in immune compromised individuals. They were not considered to be pathogenic until the outbreak of Severe Acute Respiratory Syndrome (SARS) in China and Middle East Respiratory Syndrome Coronavirus (MERS-CoV) in Middle East countries in 2002 and 2012 respectively when the viral infection led to significant deaths among the infected individuals [1]. Human coronaviruses have been shown to be rapidly evolving due to high genomic nucleotide substitution rate as well as recombination [2]. Apart from evolution, viral transmission is further fuelled by rapid urbanisation and close animal human encounters. One of the examples of pathogenicity and transmission of coronavirus is its recent outbreak in Wuhan city, China. Since the outbreak of the virus in December 2019, in Wuhan, China, people infected with this mysterious new coronavirus are escalating fast. As of 30 March 2020, more than 693000 people were infected and around 33100 people died around the globe [3]. Based on phylogenetic and taxonomy data, SARS-CoV-2 was recently shown to form a sister clade with SARS coronavirus [4]. Since SARS-CoV-2 and SARS genomes have been shown to be similar, we sought to investigate the pathogenesis of SARS-CoV-2 by predicting miRNAs regulating their pathogenesis as well as taking clues from the published study on SARS encoded non-coding RNAs [5].

MicroRNAs are known to regulate diverse biological pathways including various steps in viral replication [6]. Hence, we studied the role of host miRNAs in regulating viral replication by predicting the host miRNA binding sites among the genome of these viruses. In order to gain in depth understanding into the pathogenesis of SARS-CoV-2, we analysed all genomes of SARS-CoV-2 (n=52) that were available for download till 07 March 2020 (Supplementary figure 1). SARS reference genome was used for comparison and the analysis was performed on both the forward and reverse sequences of viral genomes. For prediction of host miRNAs that could bind to viral genomes, miRDB algorithm was used [7-8]. The analysis was performed in custom mode where user defined sequences can be uploaded and processed. All the analysis were done at default settings and only those miRNAs were studied further that had a target score >= 80. Using this approach, around 263 and 196 host miRNAs were identified that could target each of the forward and reverse SARS-CoV-2 genomic sequences (Fig1a, Supplementary table 2). Similar analysis on SARS reference genome identified 174 and 120 host miRNAs that could bind to forward and reverse sequences respectively (Fig1a, Supplementary table 2). The analysis further revealed that number of host miRNAs targeting SARS-CoV-2 genomes (both forward and reverse genomes) were significantly higher as compared to number of miRNAs targeting SARS reference genome. Despite the dissimilarity among number of host miRNAs targeting viral genomes, we observed that 86 common host miRNAs could target all the SARS-CoV-2 forward and SARS forward reference genome whereas 61 common host miRNAs were found to be targeting all the SARS-CoV-2 and SARS reverse genome (Supplementary table 3). Comparison of all the forward and reverse sequences of SARS-CoV-2 and SARS reference genome revealed 10 host miRNAs that could target all the genomes being studied and hence could be potentially used as broad acting antivirals (Fig 1b, 1c).

**Figure 1.**
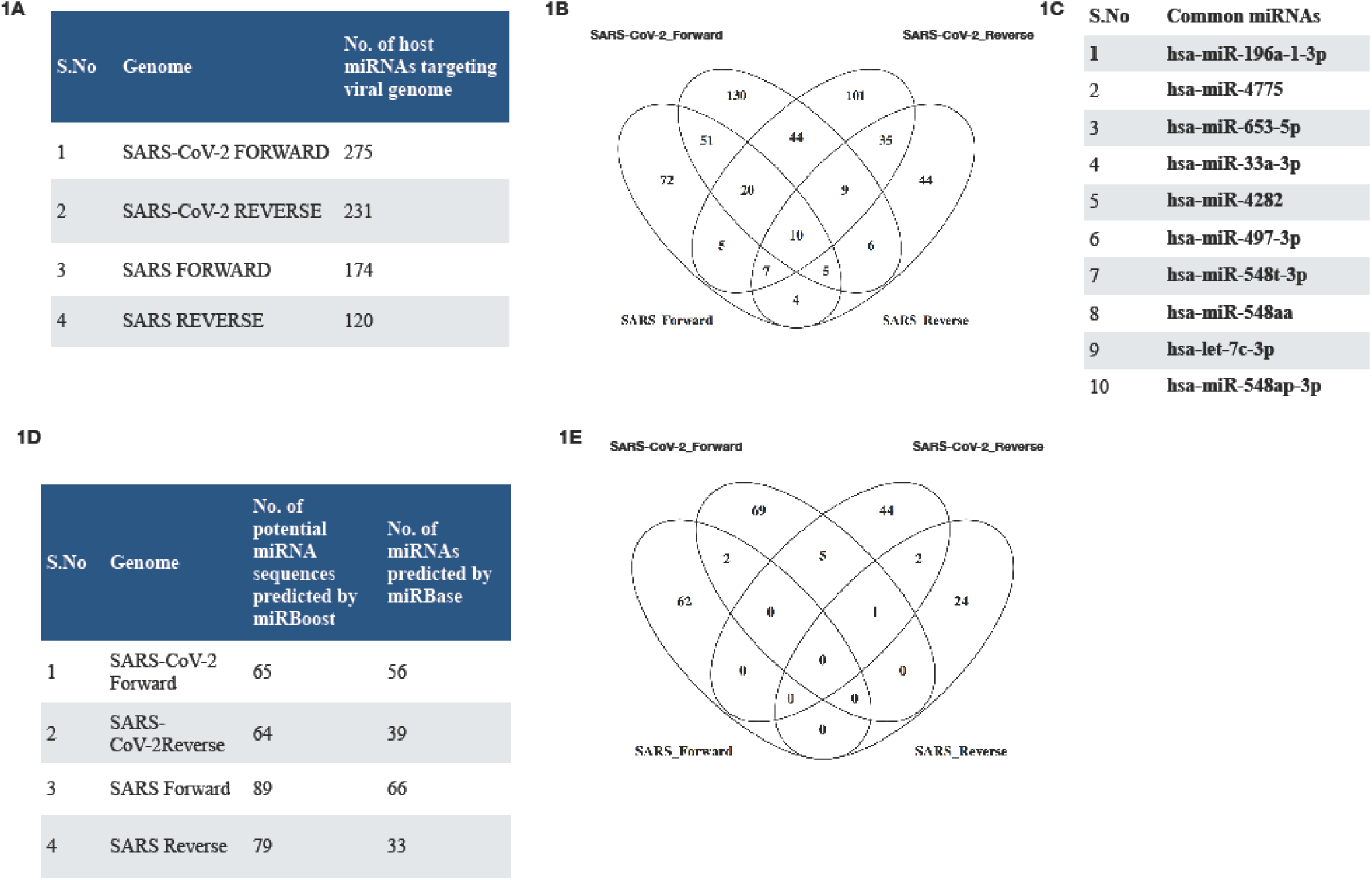
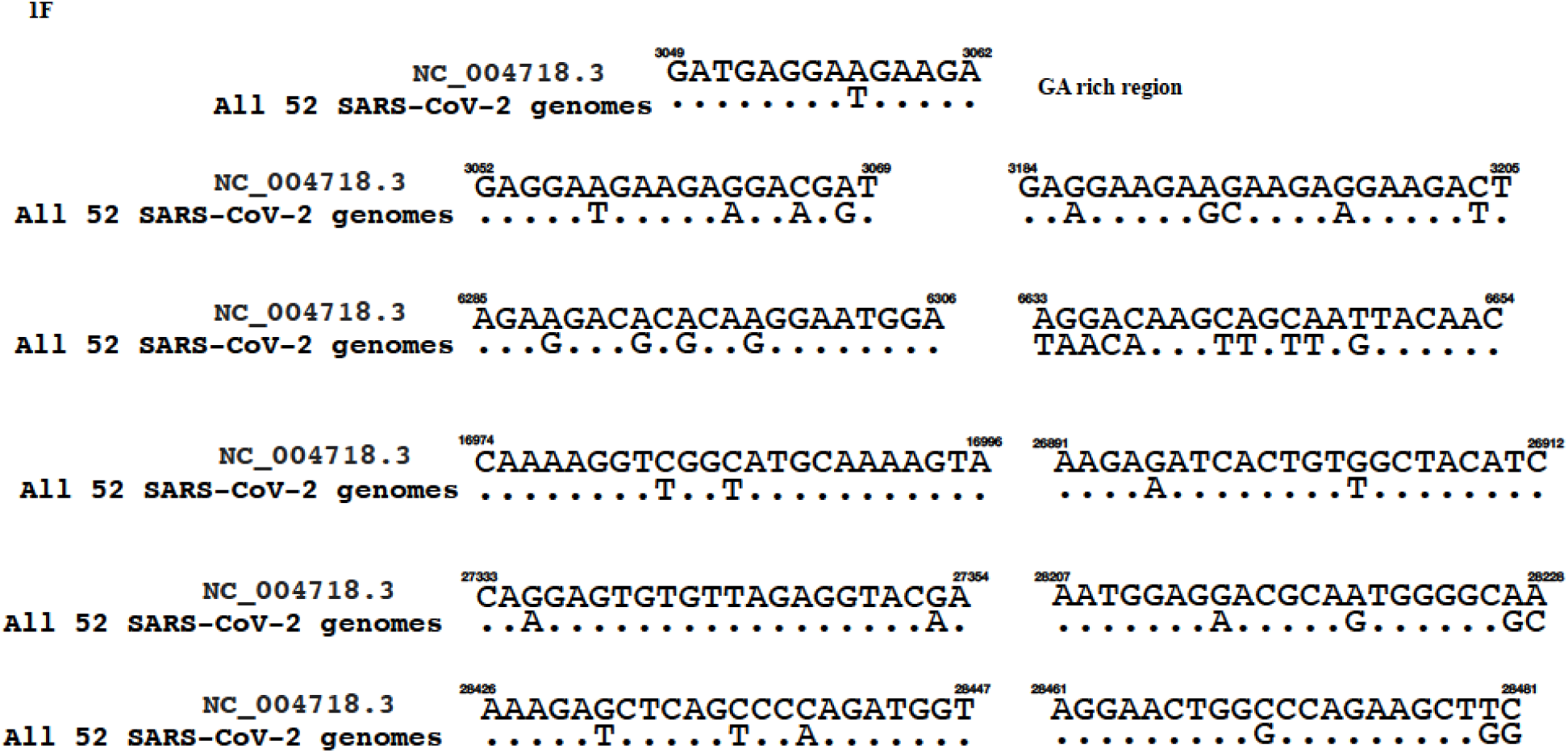
Prediction of host and viral encoded miRNAs (A) Table showing number of host miRNAs targeting various SARS-CoV-2 and SARS genomes. (B) Figure depicting various miRNAs among SARS-CoV-2 and SARS genomes. (C) Table showing the list of common miRNAs. (D) Table showing number of SARS-CoV-2 and SARS encoded miRNAs. (E) Figure depicting various common SARS-CoV-2 and SARS encoded miRNAs. (F) Figure showing comparison of SARS encoded “GA” rich region and other non-coding RNAs with that of all SARS-CoV-2 genomes.

Several viruses have been shown to regulate host signaling pathways by encoding miRNAs in their genomes. Hence, in order to gain insights into various miRNAs encoded by genomes of SARS-CoV-2 and SARS coronaviruses, a set of computational tools were used to predict miRNAs encoded by these viruses. Since hairpin loops forms the precursor of small RNAs, miRNAFold algorithm was used first to predict loops in viral genomes [9,10]. The algorithm predicted around 574 and 550 hairpin loops in each SARS-CoV-2 forward and reverse genomes respectively, 573 and 575 hairpin loops in SARS forward and reverse genome respectively. Since all hairpin loops does not result in functional miRNAs, miRBoost algorithm was then used to identify whether the hairpin structures could yield functional microRNA or not [11]. The algorithm successfully predicted around 66 miRNAs in each of SARS-CoV-2 forward genome, 64 miRNAs in each of SARS-CoV-2 reverse genome, 89 miRNAs in SARS forward genome and 79 miRNAs in SARS reverse genome. Using the above mentioned algorithms we successfully identified some of the potential viral encoded miRNAs which can be used by the viruses to modulate host signaling pathways. Then we utilized mirBase algorithm to study whether our predicted miRNAs had any similarity to human encoded miRNAs or not [12-16]. Interestingly, we observed that out of 65 sequences that were predicted by miRBoost, mirBase identified 56 miRNAs in SARS-CoV-2 forward genome sequence that matched to human specific miRNAs. Similarly out of 64 miRNAs predicted in SARS-CoV-2 reverse genome, 39 matched to human specific miRNAs. The algorithm further matched 66 and 33 miRNAs to humans from a repertoire of 89 and 79 predicted miRNAs among forward and reverse SARS genomes (fig 1d, Supplementary sheet 4). We then compared all the genomes of SARS-CoV-2 and SARS for identification of any common virus encoded miRNA. Contrary to our expectation, we did not find any virus encoded miRNA which was common to forward and reverse genomes of SARS and SARS-CoV-2 (fig 1e).

Recently, deep sequencing based approaches were used to identify SARS encoded non-coding RNAs which were then shown to play a critical role in SARS induced lung pathology [5]. The authors successfully identified a set of non-coding RNAs that were produced during SARS infections. The authors further characterized a “GA” rich region that was enriched in non-coding RNAs produced during SARS infection. Hence, we aimed to analyse SARS-CoV-2 genomic sequences for the identification of similar non-coding RNAs with a rationale that presence of similar sequences in SARS-CoV-2 might predict about the pathogenesis of the virus. Our approach revealed presence of a very similar “GA” stretch in SARS-CoV-2 which was reported in be enriched in SARS encoded non-coding RNAs. We then analyzed SARS-CoV-2 genomes for the presence of non-coding RNAs that were shown to be produced in during SARS pathology. Our comparative analysis of both viral genomes revealed that SARS-CoV-2 had very similar stretches compared to SARS encoded non-coding RNAs (figure 2). Interestingly, it was also observed that all the 52 SARS-CoV-2 genomes were highly conserved in the regions that encoded non-coding RNAs suggesting their critical role in viral pathology.

To summarize, our study identified various host and viral encoded miRNAs that could potentially regulate viral pathogenesis. We further observed that SARS-CoV-2 genomes had binding sites for more number of host miRNAs as compared to SARS genomes which might translate to low pathogenicity of SARS-CoV-2 in healthy individuals. Similarly, though SARS-CoV-2 and SARS genomes differed in terms of miRNA targets, however, there was a stark similarity among virus encoded non-coding RNAs in both the genomes thereby suggesting towards a possible overlap among pathogenesis of SARS-CoV-2 and SARS viruses and providing an opportunity to develop new therapeutics that might successfully regulate both the viruses.

## Supporting information

Supplemental Sheet 4

## Acknowledgements

VS and PA conceived the idea, designed the experiments, supervised the study wrote and finalized the manuscript. NP and SS performed the analysis. The authors declare no conflict of interest. NP is supported by UGC fellowship. VS is supported by UGC under FRP programme. The study was funded by startup grant to VS by UGC and DST PURSE grant.

## References

1. Song Zhiqi et. al. From SARS to MERS, Thrusting Coronaviruses into the Spotlight. Viruses. 2019 Jan; 11(1): 59.

2. Yvonne Xinyi Lim, Yan Ling Ng, James P. Tam, Ding Xiang Liu. Human Coronaviruses: A Review of Virus–Host Interactions. Diseases. 2016 Sep; 4(3): 26.

3. WHO situation report 70. https://www.who.int/docs/default-source/coronaviruse/situation-reports/20200330-sitrep-70-covid-19.pdf?sfvrsn=7e0fe3f8_2

4. Coronaviridae Study Group of the International Committee on Taxonomy of Viruses. The species Severe acute respiratory syndrome-related coronavirus: classifying 2019 nCoV and naming it SARS-CoV-2. Nat Microbiol. 2020 Mar 2.

5. Morales L, Oliveros JC, Fernandez-Delgado R et al. SARS-CoV Encoded Small RNAs Contribute to Infection-Associated Lung Pathology. Cell Host Microbe. 2017 Mar 8;21(3):344–355.

6. Annie Bernier, Selena M. Sagan. The Diverse Roles of microRNAs at the Host–Virus Interface. Viruses. 2018 Aug; 10(8): 440.

7. Yuhao Chen and Xiaowei Wang (2020) miRDB: an online database for prediction of functional microRNA targets. Nucleic Acids Research. 48(D1):D127–D131.

8. Weijun Liu and Xiaowei Wang (2019) Prediction of functional microRNA targets by integrative modeling of microRNA binding and target expression data. Genome Biology. 20(1):18.

9. C. Tav, S. Tempel, L. Poligny and F. Tahi. miRNAFold: a web server for fast miRNA precursor prediction in genomes. Nucleic Acids Res. 2016 Jul 8;44(W1):W181-4. S.

10. Tempel and F. Tahi. A fast ab-initio method for predicting miRNA precursors in genomes. Nucleic Acids Res. 2012 Jun;40(11):e80.

11. V.D. Tran, S. Tempel, B. Zerath, F. Zehraoui, and F. Tahi. miRBoost: Boosting support vector machines for microRNA precursor classification. RNA. 2015 May;21(5):775–85.

12. Kozomara A, Birgaoanu M, Griffiths-Jones S. miRBase: from microRNA sequences to function. Nucleic Acids Res 2019 47:D155–D162.

13. Kozomara A, Griffiths-Jones S. miRBase: annotating high confidence microRNAs using deep sequencing data. Nucleic Acids Res 2014 42:D68–D73.

14. Kozomara A, Griffiths-Jones S. miRBase: integrating microRNA annotation and deepsequencing data. Nucleic Acids Res 2011 39:D152–D157.

15. Griffiths-Jones S, Saini HK, van Dongen S, Enright AJ. miRBase: tools for microRNA genomics. Nucleic Acids Res 2008 36:D154–D158.

16. Griffiths-Jones S, Grocock RJ, van Dongen S, Bateman A, Enright AJ. miRBase: microRNA sequences, targets and gene nomenclature. Nucleic Acids Res 2006 34:D140–D144.

